# Spatial Proteomics By Parallel Accumulation-Serial Fragmentation Supported MALDI MS/MS Imaging: A First Glance Into Multiplexed and Spatial Peptide Identification

**DOI:** 10.1101/2024.11.08.622662

**Authors:** Mujia Jenny Li, Larissa Chiara Meyer, Nadine Meier, Jannik Witte, Maximilian Maldacker, Adrianna Seredynska, Julia Schueler, Oliver Schilling, Melanie Christine Föll

## Abstract

**RATIONALE:** In spatial proteomics, matrix-assisted laser desorption/ionization (MALDI) imaging enables rapid and cost-effective peptide measurements. Yet, in situ peptide identification remains challenging. Therefore, this study aims to integrate the trapped ion mobility spectrometry (TIMS)-based parallel accumulation-serial fragmentation (PASEF) into MALDI imaging of tryptic peptides to enable multiplexed MS/MS imaging.

**METHODS:** An initial MALDI TIMS MS1 survey measurement was performed, followed by a manual generation of a precursor list containing mass over charge values and ion mobility windows. Inside the dual TIMS system, submitted precursors were trapped, separately eluted by their ion mobility and analyzed in a quadrupole time-of-flight device, thereby enabling multiplexed MALDI MS/MS imaging. Finally, precursors were identified by peptide to spectrum matching.

**RESULTS:** This study presents the first multiplexed MALDI TIMS MS/MS imaging (iprm-PASEF) of tryptic peptides. Its applicability was showcased on two histomorphologically distinct tissue specimens in a 4-plex and 5-plex setup. Precursors were successfully identified by the search engine MASCOT in one single MALDI imaging experiment for each respective tissue. Peptide identifications were corroborated by liquid-chromatography tandem mass spectrometry experiments and fragment co-localization analyses.

**CONCLUSIONS:** In this study, we present a novel pipeline, based on iprm-PASEF that allows the multiplexed and spatial identification of tryptic peptides in MALDI imaging. Hence, it marks a first step towards the integration of MALDI imaging into the emerging field of spatial proteomics.

## 1 Introduction

Matrix-assisted laser desorption/ionization (MALDI) imaging is a powerful method to probe the spatial distribution of biomolecules including metabolites, lipids, peptides, and proteins. In the field of spatial proteomics, MALDI imaging stands out by enabling the antibody-independent, mass spectrometry-based, rapid, cost-effective, and spatial analysis of peptides and proteins^1–5^. Yet, in situ identification by mass spectrometry remains a major challenge in MALDI imaging since most measurements are limited to only MS1 spectra level, thus missing peptide fragment spectra (MS2), which are the backbone for peptide sequence information in mass spectrometry-based proteomics^6,7^.

As a work-around for missing MS2 spectra, MALDI imaging workflows can resort to matching MALDI-based MS1 masses to peptide identifications of additional liquid-chromatography tandem mass spectrometry (LC-MS/MS) measurements or to direct in situ MALDI MS/MS measurements that are mostly performed spot-wise^8–11^. When relying on LC-MS/MS measurements, proteome coverage is significantly deeper than in MALDI imaging. This discrepancy typically results in multiple putative identifications for a single MALDI imaging feature. As any LC-MS/MS precursor within an adequate mass accuracy window has to be considered correct, the probability of incorrect peptide identification and biological interpretation is increased, highlighting the need for defining specific matching thresholds^6,12,13^.

Direct in situ identification of peptide ions in MALDI imaging, i.e. by ion fragmentation, is considered optimal but is hampered by several hurdles^7,14,15^: First, the lack of chromatographic separation (as used for LC-MS/MS) leads to the simultaneous detection of near-isobaric peptides resulting in co-fragmentation and chimeric MS2 spectra as well as limited enrichment of peptide ions. Second, most precursor selection methods for MALDI imaging typically filter for one precursor ion per set of laser shots, thus hindering multiplexing capabilities, limiting precursor selection windows and resulting in higher sample consumption and longer measurement times.

During recent years, quadrupole time-of-flight (Q-TOF) tandem mass spectrometry has been enhanced by preceding trapped ion mobility spectrometry (TIMS) for both electrospray ionization (ESI) and MALDI systems. A dual TIMS cell system sorts and traps precursor ions based on their collisional cross section (CCS), thereby allowing separation of isobaric and isomeric peptides. The TIMS cell measures the closely related ion mobility (*K*) value. Within this study, we report the inverse reduced ion mobility (1/*K*_0_) which standardizes the ion mobility for temperature and pressure.

TIMS-based synchronization of ion storage and release with subsequent, quadrupole-based ion selection and fragmentation steps leads to very efficient ion usage; a setup called parallel accumulation-serial fragmentation (PASEF)^16^. This method has already significantly advanced ESI systems as used in LC-MS/MS shotgun proteomics and was also applied to targeted proteomics as parallel reaction monitoring (prm)-PASEF ^17–23^ More recently, novel acquisition software enhancements have made the prm-PASEF mode available to commercial MALDI imaging systems such as in the timsTOF fleX (Bruker Daltonics, Bremen, Germany), thereby enabling the multiplexed identification of peptide ions per set of laser shots. This software tool is called iprm-PASEF (Bruker Daltonics, Bremen, Germany) and its applicability has already been demonstrated for lipids and metabolites^24,25^.

The present study is the first to optimize and apply TIMS-based acquisition modes including iprm-PASEF to MALDI imaging of tryptic peptides for improving peptide identifications.

## 2 Materials and Methods

### 2.1 Tissue Specimens and Tissue Processing

Mouse kidney tissues were obtained as surplus tissue from animals that had been sacrificed for unrelated experimental purposes in accordance with approved animal experimentation protocols. The mouse kidney samples represent leftover material that would have been discarded otherwise. Breast cancer tissue from murine patient-derived xenografts (PDX tumor) were provided by Charles River Laboratories, Freiburg, Germany. The PDX experiments were carried out in strict accordance with the recommendations in the Guide for the Care and Use of Laboratory Animals of the Society of Laboratory Animals (GV SOLAS). The PDX experiments were approved by the Committee on the Ethics of Animal Experiments of the regional council (Permit Numbers: I19-02). Bovine liver was obtained commercially from a local butcher. All tissues were formalin-fixed and embedded in paraffin (FFPE) according to standard protocols^26^. FFPE blocks were sliced into 2 µm thick sections and mounted onto IntelliSlides (Bruker Daltonics, Bremen, Germany) or indium tin oxide (ITO) slide (Bruker Daltonics, Bremen, Germany).

### 2.2 Peptide Samples

For standard peptides and iprm-PASEF validation, the synthetic peptide iRT kit was used (Biognosys AG, Schlieren, Switzerland). For LC-ESI-TIMS-MS/MS optimization, Pierce HeLa Protein Digest Standard (Thermo Fisher Scientific, Waltham, Massachusetts, USA) was used. Further synthetic peptides were used for iprm-PASEF validation: Apelin-13 (Santa Cruz Biotechnology, Dallas, USA), Substance P (Merck, Darmstadt, Germany), and two further synthetic peptides (KLKESYCQRQGVPMN, KLKVIGQDSSEIHFKV) (GenScript, Piscataway Township, New Jersey, USA).

### 2.3 Sample Preparation for MALDI Imaging of Tryptic Peptides

For all MALDI imaging measurements, two adjacent tissue slides were prepared simultaneously as described previously^27^. Deparaffinization was carried out in a series of xylol and ethanol/water dilutions as described previously^26^. Tissue sections were washed twice with 10 mM ammonium bicarbonate (AmBC) for 1 min. Antigen retrieval was performed in a citric acid monohydrate buffer with pH 6.0 in a steamer for 1 h at approximately 100 °C. Slides were rinsed twice with 10 mM AmBC and dried for 10 minutes in a vacuum atmosphere at room temperature. Freshly prepared N-tosyl-L-phenylalanine chloromethyl ketone treated trypsin (Worthington, Lakewood, NJ, USA) (0.1 mg/mL in 10 % acetonitrile (ACN), 40 mM AmBC) was sprayed on the tissue slides using the M3+ sprayer (HTX Technologies, Chapel Hill, NC, USA) with the following parameters: Temperature 30 °C, nozzle velocity 750 mm/min, flow rate 30 μL/min, number of passes 8, track spacing 2 mm, pattern CC, drying time 0 s, and nitrogen gas pressure set to 10 psi. Afterwards, slides were incubated in a humidified chamber for 2 h at 50 °C. Alpha-cyano-4-hydroxycinnamic acid matrix (CHCA) (Sigma-Aldrich, Munich, Germany) (10 mg/mL, 70 % ACN, 0.1 % trifluoroacetic acid (TFA), 7 mM ammonium phosphate) was applied using the M3+ sprayer with the following parameters: Temperature 60 °C, nozzle velocity 1350 mm/min, flow rate 100 μL/min, number of passes 8, track spacing 3 mm, pattern CC, drying time 10 s, and nitrogen gas pressure set to 10 psi. All samples within one experiment were stored for the same time and measured consecutively and within 24 hours upon sample preparation. For validation purposes, two additional setups were carried out: First, for section 3.2, FFPE bovine liver slides were prepared for MALDI imaging measurement in triplicates as described above. The iRT kit peptides were spotted twice in a dilution series (400 fmol, 200 fmol, 100 fmol, 50 fmol in 0.5 µL droplets) on top of the digested liver section before matrix application. Second, for section 3.3, 1 µL of a mastermix containing iRT kit peptides, Apelin-13, Substance P, and synthetic peptides (KLKESYCQRQGVPMN, KLKVIGQDSSEIHFKV) were mixed with 1 µL CHCA matrix and spotted onto an empty ITO slide in triplicates to compare 1-plex and 5-plex iprm-PASEF measurements. The mastermix contained the following concentrations: iRT kit peptides (566.37 fmol/uL), Apelin-13 (0.0088 µg/µL), Substance P (0.0177 µg/µL), KLKESYCQRQGVPMN (0.221 µg/µL) and KLKVIGQDSSEIHFKV (0.044 µg/µL).

### 2.4 MALDI TIMS MS1 Imaging Acquisition

MALDI TIMS MS1 imaging datasets were acquired in the positive-ion reflector mode using a timsTOF fleX mass spectrometer (Bruker Daltonics, Bremen, Germany) at a pixel size of 50×50 μm, using 600 shots and a laser frequency of 10 kHz. The instrument is equipped with a Nd:YAG laser emitting at 355 nm. The *m*/*z* range was set to *m*/*z* 800-2,000, the 1/*K*_0_ range to 1.2-2.1 V•s/cm^2^ with a ramp time of 230 ms.

Tuning parameters were optimized and set as follows: MALDI plate offset 80 V, deflection 1 delta 80 V, funnel 1 RF 250 Vpp, funnel 2 RF 500 Vpp, multiple RF 500 Vpp, collision energy 10 eV, collision RF 4000 Vpp, ion energy 5 eV, transfer time 185 ms, and pre pulse storage 20 µs. The TIMS cell pressure was decreased by adjusting the nitrogen flow until the calibration mass *m*/*z* 1221 (Hexakis(1H, 1H, 4H-hexafluorobutyloxy)phosphazine (Agilent Technologies, Santa Clara, USA)), aligned to 170 V, yielding a TIMS cell pressure of 2.0 mbar. 1/*K*_0_ and *m*/*z* calibration was performed over ESI with the ESI-low concentration tuning mix (Agilent Technologies, Santa Clara, USA). For validation purposes, MALDI TIMS off MS1 imaging measurements were carried out with TIMS turned off and all other parameters kept identical.

### 2.5 MALDI TIMS MS1 Imaging Data Analysis: Precursor Selection by Statistical Analysis

For precursor selection, MALDI TIMS MS1 imaging data was analyzed with the software SCiLS Lab (Version 2024a Core, Bruker Daltonics, Bremen, Germany). The variance was kept to 10 ppm for *m*/*z* and 0.01 V•s/cm^2^ for 1/*K*_0_. Feature finding was carried out with T-ReX^3^ (TIMS) algorithm, root mean square (RMS) normalization, 2×2 spatial smoothing, 50 % coverage and 1 % relative intensity threshold. In cases where feature selection was slightly off, the peak maximum was manually selected in the two-dimensional *m*/*z* and 1/*K*_0_ plot. For the statistical analysis, the “Segmentation” tool in SCiLS Lab was used with RMS normalization, the region set to the tissue region, T-ReX^3^ (TIMS) feature list, denoising set to none, “Bisecting k-Means” as method and “Correlation Distance” as metric. For the “Find Values Co-Localized to Region” tool, RMS normalization was selected, ion images of the entire tissue region and the T-ReX^3^ (TIMS) feature list were submitted and correlated with one of the segmented regions of interest determined by the spatial segmentation tool. The co-localization algorithm uses Pearson correlation analysis and only considers statistically significant correlations with p less than 0.05.

### 2.6 Sample Preparation of Peptides Co-Crystallized with MALDI Matrix for LC-ESI-TIMS-MS/MS

To recover the co-crystallized peptides after MALDI imaging acquisition, the matrix layer on the tissue slide was dissolved after MALDI measurement, collected with 70 % ACN, and vacuum-dried for ACN removal^12^. Peptides were cleaned up with PreOmics Cartridges (PreOmics, Planegg/Martinsried, Germany). The eluted peptides were vacuum-dried and dissolved in water and 0.1 % formic acid before loading on Evotips (Evosep, Odense, Denmark) for final desalting and LC-ESI-TIMS-MS/MS measurement.

### 2.7 LC-ESI-TIMS-MS/MS Acquisition Focused on Singly Charged Peptides

LC-ESI-TIMS-MS/MS measurements of co-crystallized peptides and 100 ng HeLa digest for method evaluation were conducted on a timsTOF fleX mass spectrometer equipped with a CaptiveSpray ion source (Bruker Daltonics, Bremen, Germany). 10 fmol iRT Kit was spiked into all samples before Evotip loading. Chromatographic separation was performed with the Evosep One HPLC system (Evosep, Odense, Denmark), using the 30 samples per day method (44 min gradient, 500 nL/min) and the EV1137 performance 15 cm reversed phase column (Evosep, Odense, Denmark) with buffer A consisting of 0.1 % aqueous formic acid and buffer B of 0.1 % formic acid in ACN. Samples were measured in positive data-dependent acquisition (DDA)-PASEF mode. To increase measurement of singly charged peptides, adjustment of the TIMS cell pressure and ion mobility calibration were performed as described in MALDI TIMS MS1 acquisition. The mass range was set to *m/z* 100–1,700 and 1/*K*_0_ range to 0.7–2.1 V•s/cm^2^. Collision-induced dissociation energy was set to 20–75 eV, ion accumulation time to 166 ms with a 100 % duty cycle. The PASEF precursor polygon region was adjusted manually to include singly charged peptide traces. Tuning parameters were optimized and set as follows: Deflection 1 delta 80 V, funnel 1 RF 250 Vpp, funnel 2 RF 500 Vpp, multiple RF 500 Vpp, collision energy 10 eV, collision RF 2,500 Vpp, ion energy 5 eV, transfer time 70 ms, and pre pulse storage 15 µs. The capillary voltage was set to 1,600 V and the drying gas flow rate was 3 L/min with 180 °C drying temperature.

### 2.8 LC-ESI-TIMS-MS/MS Data Analysis

The FragPipe pipeline (v22.0) was employed for the analysis of the DDA-PASEF datasets^28–30^. Peptide identification of MS2 spectra was obtained using MSFragger with raw (.d) files as input. Protein databases, either human (UP000005640, downloaded August 13, 2024) or mouse (UP000000589, downloaded June 10, 2024), were both compiled with common contaminants, and internal standards (iRT Kit peptides). Reversed sequences were added as decoys. Default settings for label free quantification were applied with proline +15.999 (hydroxylation) set as a variable modification in MSFragger.

### 2.9 iprm-PASEF Acquisition

The iprm-PASEF datasets were acquired using a prototype of the iprm-PASEF tool for timsControl 5.1 provided by Bruker Daltonics GmbH, Bremen, Germany. Measurements were performed in MS/MS and positive ion reflector mode on the timsTOF fleX at a pixel size of 100×100 μm, using 2,000 shots and a laser frequency of 5 kHz. The *m*/*z* range was set to *m*/*z* 100–2,000, the 1/*K*_0_ range was set to 1.2–2.1 V•s/cm^2^ with a ramp time of 230 ms. Tuning parameters were optimized and set as follows: Collision RF 750 Vpp, ion energy 5 eV, transfer time 55 ms, pre pulse storage 8 µs. Adjustment of the TIMS cell pressure and 1/*K*_0_ calibration were performed as described in MALDI TIMS MS1 imaging acquisition. The iprm-PASEF methods were set up with predefined 1/*K*_0_ and *m*/*z* ranges. Collision energies were adjusted depending on the precursor mass from 45–140 eV. For validation purposes, MALDI TIMS off MS/MS imaging measurements were performed with only TIMS turned off and all other parameters kept identical.

### 2.10 iprm-PASEF Data Analysis

In the DataAnalysis software (Bruker Daltonics, Bremen, Germany), MS2 spectra of each individual precursor were extracted over the mobilogram, averaged over all pixels, exported from the iprm-PASEF datasets as individual “.mgf” files and submitted to the “MASCOT MS/MS Ions Search” (Matrix Science, London, UK) against the Swiss-Prot (2024/04) database including contaminants filtered by the respective taxonomy (mus musculus, homo sapiens)^31^. Enzymatic digestion was set to “Trypsin” with two missed cleavages allowed. The precursor tolerance was set to 100 ppm, fragment ion tolerances to 0.3 Da, and target FDR to 1 %. Histidine, tryptophane, proline and methionine oxidation as variable modification were included. MS2 spectra and assigned theoretical b- and y-ions were visualized using the pyteomics package in python^32^. Theoretical b- and y-ions were matched with a tolerance of 0.3 Da. Ion images of peptide fragments were plotted in SCiLS Lab. For co-localization analysis of fragments, features were picked manually in the two-dimensional *m*/z and 1/*K*_0_ plot in SCiLS Lab. The obtained feature list was then submitted to the “Find Values Co-Localized to Feature” tool, normalization was set to RMS, images of the tissue region were selected and the parent ion as feature to correlate. The co-localization algorithm uses Pearson correlation analysis and only considers statistically significant correlations with p less than 0.05. Finally, co-localization scores were obtained to statistically estimate the spatial correlation of fragment to parent ion images.

## 3 Results

### 3.1 Overview of PASEF-Enabled MS/MS MALDI Imaging of Peptides

Spatial proteomics using MALDI imaging benefits from the direct, in situ MS/MS based identification of peptide ions. PASEF-enabled MALDI imaging supports this approach by ion mobility-based precursor separation and entrapment according to distinct ion mobility features, thereby allowing multiplexing per set of laser shots. For measurement in iprm-PASEF, precursor ions of interest are selected within predefined ranges of their relevant molecular properties, namely *m*/*z* and 1/*K*_0_. The spatial proteomics pipeline performed in this study consists of four steps: (A) An initial MALDI TIMS MS1 survey measurement (50×50 µm), (B) manual assembly of a precursor list, (C) iprm-PASEF based MALDI imaging for the creation of MS2 spectra of those precursors (100×100 µm), and (D) a peptide to spectrum matching (PSM) and potential corroboration. We used 50 µm resolution in the MS1 scan for refined spatial analysis, while iprm-PASEF at 100 µm for identification focused on a faster acquisition and higher spectral quality.

The initial MALDI TIMS MS1 measurement (Figure 1A) yields an overview of locally abundant MS1 precursor ions including *m*/*z* and 1/*K*_0_ values from which a feature list (typically 2,000-8,000 features) is derived. Multiplexed precursor ion trapping in the first TIMS chamber for iprm-PASEF relies on non-overlapping 1/*K*_0_ windows. In addition, those windows need to be sufficiently wide to allow for efficient isolation. These requirements typically limit the number of precursor ions to 25 entries with distinct ion mobility windows. Therefore, a sophisticated precursor selection is necessary to perform the subsequent iprm-PASEF survey.

**Figure 1:**
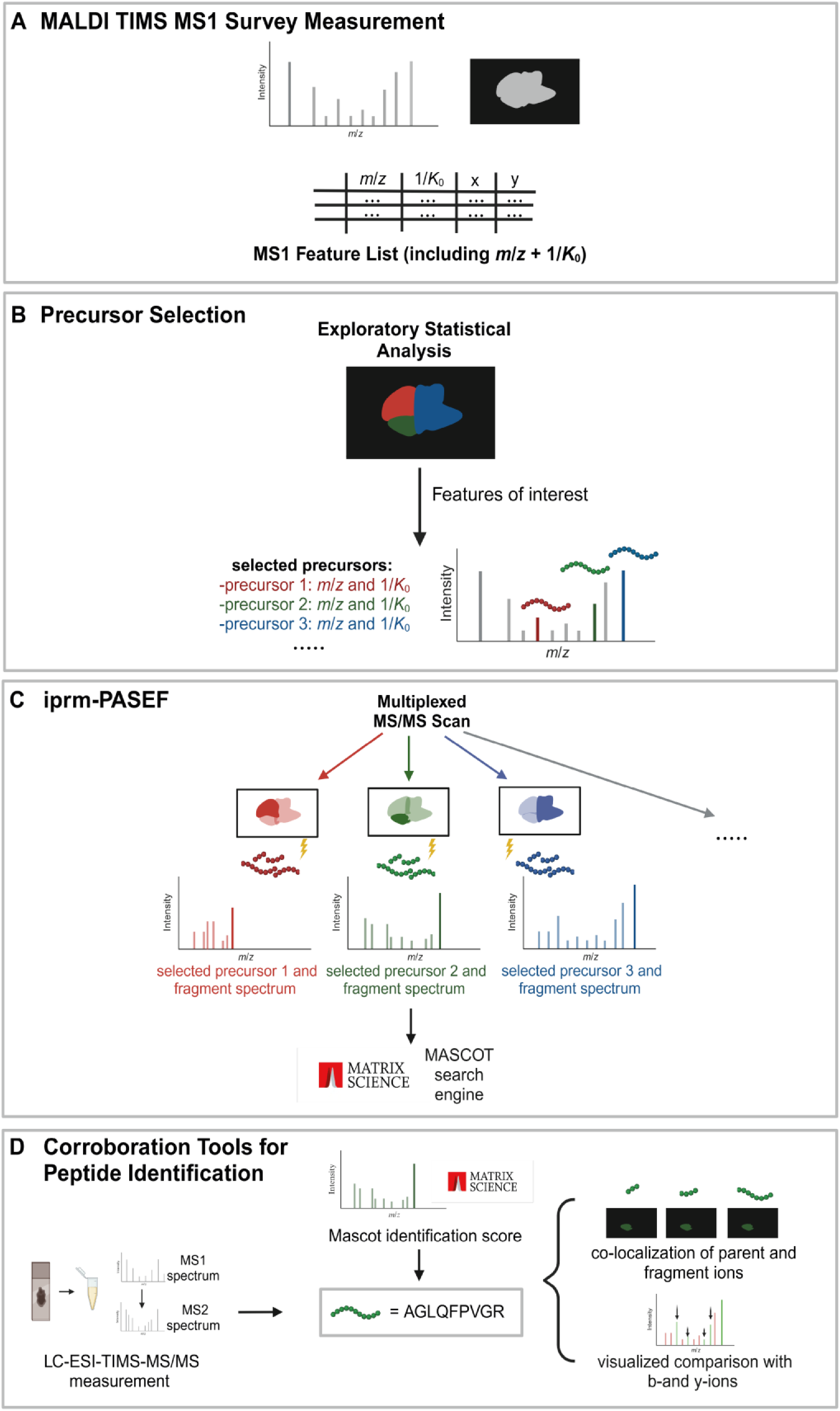
Overview of the iprm-PASEF pipeline presented in this study. **(A)** TheMALDI TIMSMS1survey imaging scan provides a feature list. **(B)** Precursor selection, e.g. by an exploratory statistical analysis, is performed for the targeted MS/MS measurement. **(C)** With the selected precursors and their distinct *m/z* and *1/K_0_* values, a multiplexed MALDI TIMS MS/MS imaging (iprm-PASEF) scan is performed. The resulting MS2 spectra are identified using the MASCOT search engine. **(D)** Overview of the corroboration tools used in this study. (Created in BioRender. Schilling, 0. (2024) https://BioRender.com/g48n665).

As iprm-PASEF is a targeted workflow, the precursor definition relies on manual selection (Figure 1B). Notably, this selection highly depends on the experimental design and can be individually tailored to the aim of the study. For example, a statistical analysis of the MALDI TIMS MS1 dataset including a segmentation/classification and co-localization can reveal features of histomorphological interest that subsequently can be chosen for further fragmentation. Eventually, a customized precursor list containing *m*/*z* values and 1/*K*_0_ windows is obtained from the MALDI TIMS MS1 imaging dataset.

The iprm-PASEF proof-of-concept measurement is performed at 100×100 µm spatial resolution, resulting in measurement times less than 100 min for 1 cm^2^ area (2,000 number of laser shots with 5 kHz frequency). In our preliminary experience, the reduction to 5 kHz enables prolonged accumulation times that benefit spectra intensity. The entire tissue section is scanned in iprm-PASEF mode resulting in spatially resolved information of fragment ions. For subsequent PSM of individual MS2 spectra, we resorted to the online version of the well-established MASCOT software. For this purpose, ion mobility filtered MS2 spectra are averaged and exported to the “.mgf” format as implemented in the DataAnalysis software (Figure 1C).

The presented pipeline further includes additional analyses to corroborate peptide identification by iprm-PASEF (Figure 1D): To complement the online MASCOT search output that is restricted to solely scores, mean MS2 spectra and correctly detected b- and y-ions of the identified peptides are visualized with the pyteomics package. Ion images of fragments are examined to show similar spatial distribution as expected and statistically analyzed by the “Find Values Co-Localized to Feature” tool in SCiLS Lab. With this tool, co-localization of fragment to parent ions is evaluated based on Pearson correlation. We further aimed to support the iprm-PASEF peptide identifications with PSM of LC-ESI-MS/MS (incl. TIMS) data. This step can be pursued with additional sample material or with matrix extracted peptides that have remained following initial MALDI imaging^12^. If the same peptide is identified by LC-ESI-TIMS-MS/MS, precursor *m*/*z* and 1/*K*_0_ values can be compared with MALDI imaging data to corroborate their identities.

### 3.2 Inclusion of Dual TIMS into MALDI MS1 Imaging

As outlined, usage of TIMS for selection and entrapment of precursor ions is fundamental for iprm-PASEF. We aimed to probe whether the dual TIMS step prior to Q-TOF measurement impacts MS1 sensitivity as compared to a Q-TOF measurement on its own. To this end, we adjusted TIMS parameters to accommodate for peptidic precursors: The ramp time was increased to 230 ms for sufficient separation and the TIMS cell nitrogen gas flow was lowered to 2.0 mbar to optimize accumulation.

To evaluate the effect of upstream dual TIMS usage on MS1 sensitivity, the iRT peptide kit was spotted in a dilution series on trypsin-digested bovine liver tissue using default settings for MALDI MS1 imaging. Without dual TIMS (hence Q-TOF on its own), the lowest detectable spot was at 50 fmol per peptide. Inclusion of the dual TIMS step for MALDI TIMS MS1 yielded a very comparable result (Figure 2A). For all iRT ion images see Supplementary Figure 1. We conclude that inclusion of the dual TIMS step for MALDI MS1 does not significantly affect MS1 sensitivity.

**Figure 2:**
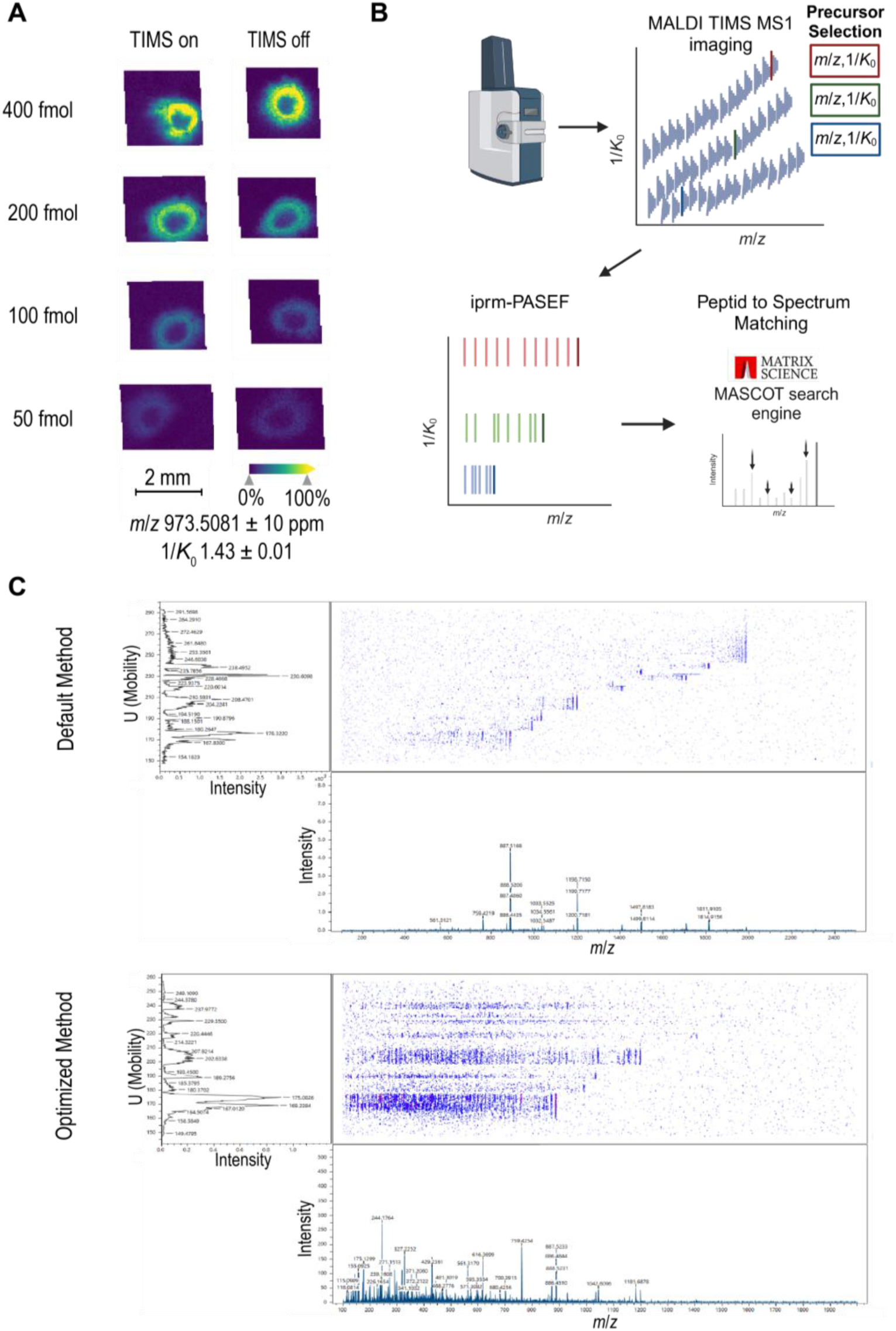
Evaluation of TIMS-supported MALDI MS1 and MS/MS imaging of tryptic peptides. **(A)** Synthetic tryptic peptides (iRT kit) were spotted on a bovine slice tissue section, CHCA-matrix was applied and MALDI MS1 imaging measurement was carried out either in TIMS on or TIMS off mode. The data was visualized and analyzed using the SCiLS Lab software. (B) Overview of the iprm-PASEF methodology. After a MALDI TIMS MS1 imaging scan, specific precursors are selected for a MALDI TIMS MS/MS imaging (iprm-PASEF) scan. MS2 spectra are analyzed using MASCOT. (Created in BioRender. Schilling, 0. (2024) https://BioRender.com/j85k835) (C) Comparison of the mobility/ *mlz* heatmap between the default and optimized iprm-PASEF method visualized in timsControl (Broker Daltonics, Bremen, Germany).

### 3.3 iprm-PASEF of Tryptic Peptides

iprm-PASEF is an implementation of multiplexed MALDI TIMS MS/MS imaging to generate spatially resolved MS2 spectra (Figure 2B). When establishing the TIMS-enabled, multiplexed MS/MS data acquisition of tryptic peptides, we noticed the need to optimize the acquisition method to enable the detection of low *m/z* fragment signals with additional sensitivity for high molecular weight precursor masses (Figure 2C). Compared to the default peptide imaging method provided by Bruker Daltonics, we adjusted relevant tuning parameters as follows: “Collision RF” from 4,000 Vpp to 750 Vpp, “Transfer Time” from 185 ms to 55 ms and “Pre Pulse Storage” from 20 µs to 8 µs. The *m*/*z* range was set to *m*/*z* 100-2,000. Collision energies were increased dependent on *m*/*z* from 45 to 140 eV.

We further evaluated the impact of dual TIMS on MS2 spectra quality. Equally sized areas of tryptic digested bovine liver tissue slice were imaged in MS/MS mode in triplicates. Three precursors were selected for the MS/MS measurement (Supplementary Table 1). Apart from TIMS turned on or off, all other parameters were kept identical. Since independent measurements are required for every single precursor with TIMS turned off, the measurement area was tripled and measurements took longer compared to iprm-PASEF. All MS2 spectra per precursor were averaged, exported in the “.mgf” format, and submitted to MASCOT for PSM. In iprm-PASEF, all precursors were matched to peptide sequences with MASCOT scores higher than 35. Without the TIMS dimension, only one precursor was matched to a peptide sequence. However, for TIMS off methods, individual, precursor-specific refinement of acquisition parameters may yet improve spectral quality and was not pursued in the present study. Overall, iprm-PASEF allows for the simultaneous and multiplexed acquisition of MS2 spectra over a wide precursor *m*/*z* range (Figure 3).

**Figure 3:**
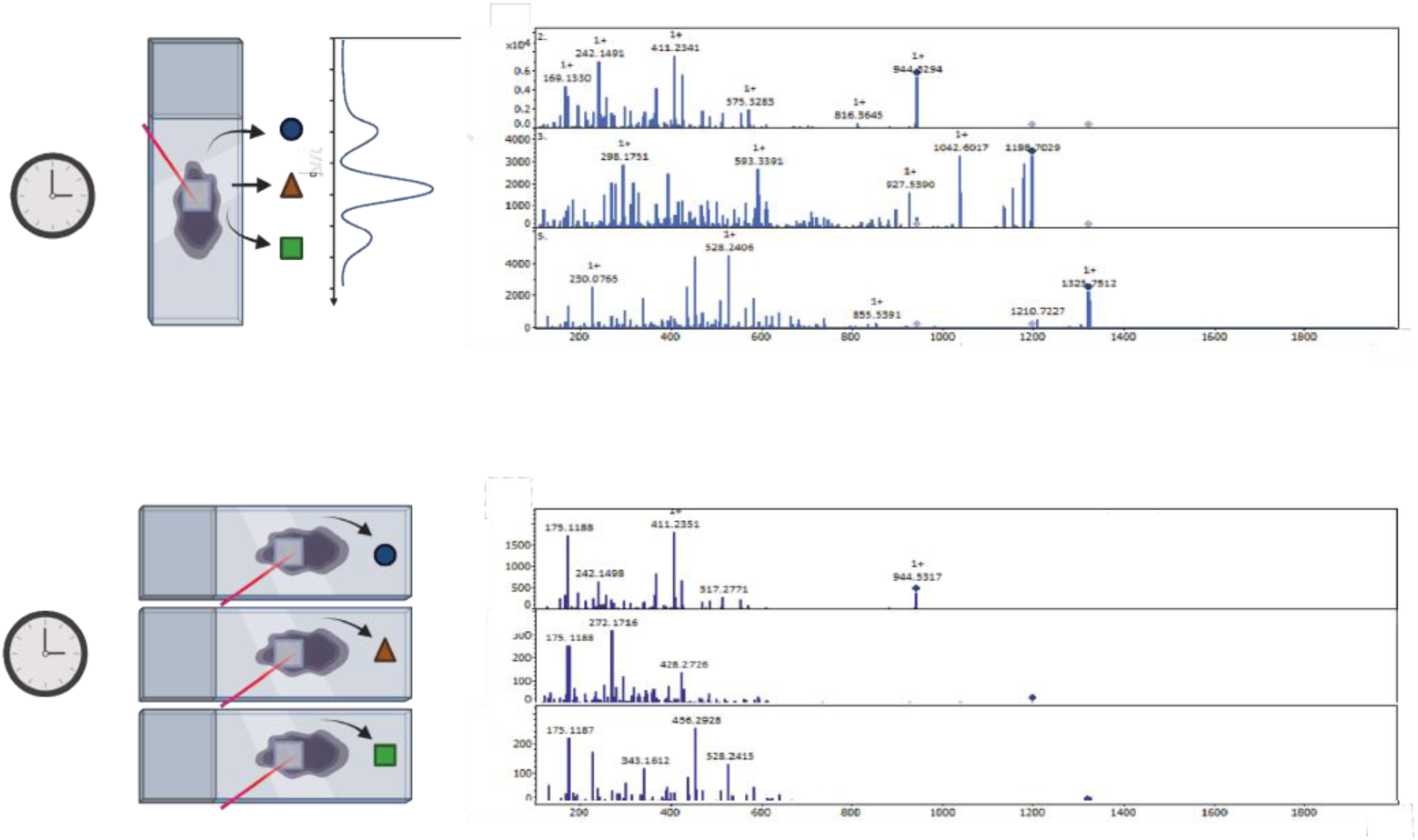
Validation of the multiplexed MALDI MS/MS imaging using the novel iprm-PASEF Software. Comparison between iprm-PASEF (TIMS on) and MALDI TIMS off MS/MS imaging measurements. Three precursors (m/z 944.5317, *mlz* 1198.7029 and m/z 1325.7512) are targeted in both modes on a prepared bovine liver tissue slice. In iprm-PASEF, all three precursorscan be measured simultaneously whereas in MALDI MS/MS imaging in TIMS off mode only one precursor can be targeted at a time. If possible, precursors were identified using MASCOT. Mass accuracies obtained by MASCOT are shown in ppm. (Created in BioRender. Schilling, 0. (2024) https://BioRender.com/j51r217, modified)

Next, we assessed whether multiplexing in iprm-PASEF affects MS2 spectrum quality. A mastermix of five synthetic peptides (see Supplementary Figure 2 for sequences) was mixed with CHCA matrix and spotted directly on an empty ITO slide (see section 2.2). Then, iprm-PASEF measurements were performed; either using multiplexing of all five precursors or using 1-plex, thereby individually aiming for one precursor after another. Visual inspection of the resulting MS2 spectra suggests highly similar MS2 spectra from both measurement approaches (Supplementary Figure 2). For further comparison of MS2 spectra quality, the completeness of the expected b- and y-ion series was determined and can be found in the Supplementary Table 2. Notably, the number of detected b- and y-ions showed a maximum variance of two ions between measurements. We therefore conclude that multiplexing iprm-PASEF measurements does not affect MS2 spectra quality.

### 3.4 Optimization of LC-ESI-TIMS-MS/MS Measurement to Focus on Singly Charged Peptides

LC-ESI-TIMS-MS/MS based peptide identifications may serve to corroborate MALDI TIMS MS/MS based peptide identifications. However, MALDI ionization produces mostly singly charged ions in the typical mass range of tryptic peptide, whereas ESI usually yields tryptic peptide ions charged 1+ to 4+. For improved comparability of MALDI and ESI data, we sought to focus on singly charged ions in LC-ESI-TIMS-MS/MS since 1/*K*_0_ values are charge state dependent^33–35^. Therefore, we optimized the tuning parameters in LC-ESI-TIMS-MS/MS measurements to increase the detection of singly charged peptides. An overview of an optimization experiment testing different settings can be found in Supplementary Table 3. Key adjustments include setting the TIMS cell pressure similar to the MALDI imaging acquisition, increasing the upper 1/*K*_0_ limit and redefining the PASEF polygon precursor region to include singly charged precursor ions.

Next, we assessed whether 1/*K*_0_ values for the same peptide sequence are comparable between LC-ESI-TIMS-MS/MS and MALDI TIMS MS1 imaging. To this end, iRT peptides (Biognosys AG, Schlieren, Switzerland) were measured using LC-ESI-TIMS-MS/MS and MALDI TIMS MS1 imaging. The *m*/*z* and 1/*K*_0_ values were highly comparable between those two measurements with an *m*/*z* variance of lower than 5 ppm and a 1/*K*_0_ variance of lower than 0.01 V•s/cm^2^ (Table 1).

**Table 1:**
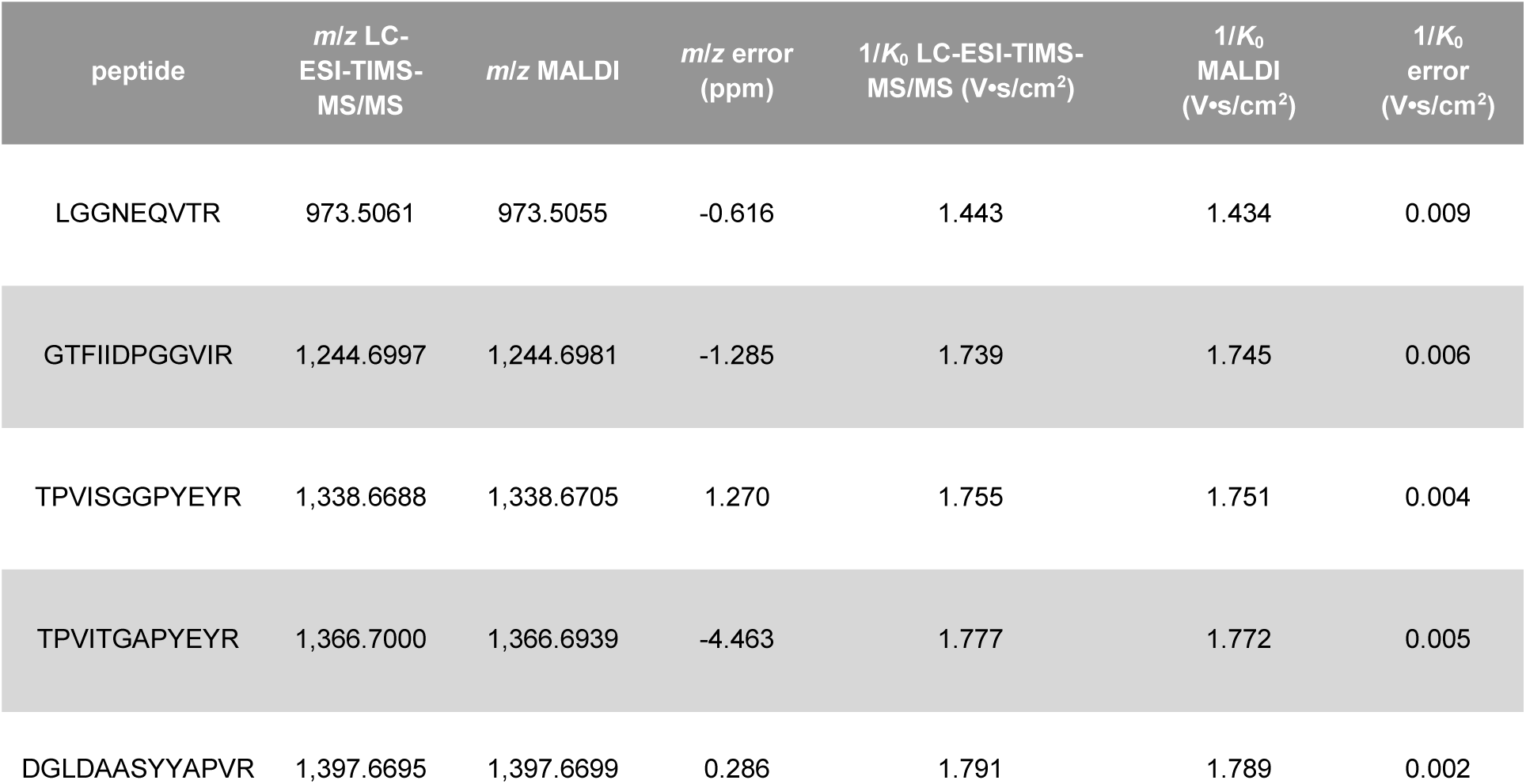
*m*/*z* and 1/*K*0 values of iRT peptides obtained from MALDI imaging spots on bovine liver. In LC-ESI-TIMS-MS/MS, 10 fmol of iRT Kit was measured together with 100 ng HeLa digest.

### 3.5 Exemplary Application to FFPE Tissue Specimens

To further elaborate the novel, MS/MS-enabled, MALDI-based spatial proteomics technique, two samples were selected as histomorphologically distinct tissue specimens. An FFPE mouse kidney sample was analyzed in a 5-plex setting and breast cancer tissue from murine patient-derived xenografts (PDX tumor) was measured in a 4-plex setup, demonstrating the applicability of the workflow to different tissue species.

#### 3.5.1 MALDI TIMS MS1 Survey Measurement

MS1 peptide imaging took 138 minutes generating 30,245 pixels (PDX tumor) and 107 minutes generating 22,977 pixels (mouse kidney) with a spatial resolution of 50×50 µm, resulting in a measurement speed of approximately 3.5 pixels per second, which is less than 200 min per cm^2^ of tissue. Initial data assessment including feature finding, data visualization and normalization was carried out in SCiLS Lab. Feature finding using T-ReX^3^ (TIMS) algorithm resulted in 6,119 (mouse kidney) and 7,536 (PDX tumor) features, respectively.

#### 3.5.2 Precursor Selection by Statistical Analysis

As mentioned above, precursor selection for iprm-PASEF, a targeted method, is highly dependent on the experimental goal. This study focuses on method development and aims to provide a first proof of concept for spatial and multiplexed peptide identification. Therefore, we resorted to a basic exploratory statistical analysis of the MALDI TIMS MS1 imaging dataset. First, a “Bisecting k-Means” segmentation was performed in SCiLS Lab with the T-ReX^3^ (TIMS) feature finding output to separate the two most distinct tissue subregions (Figure 4A, red and green). These two subregions correspond to the extracellular space (green segment) and the cellular compartment (red segment), respectively. The tool “Find Values Co-Localized to Region” in SCiLS Lab was then applied to all ion images of the T-ReX^3^ (TIMS) feature list, with the green subregion selected as region of interest. Thresholds of higher than 0.7 or lower than - 0.7 were applied. Accordingly, scores higher than 0.7 indicate a strong co-localization with the green subregion and scores lower than −0.7 with the red subregion, respectively. The results were further combined with a literature search in PubMed to obtain putative peptide identifications by *m*/*z* value. We included this potentially biased refinement step to increase confidence in the correctness of peptide identifications, as this study is the first to multiplex peptide fragmentation and serves as a proof of concept. Finally, the precursor selection for the PDX tumor included *m*/*z* 898.5029 and *m*/*z* 1,443.6980 with high co-localization scores to extracellular space (0.74 and 0.76), and *m*/*z* 944.5373 and *m*/*z* 1,325.7528 with low co-localization scores (−0.71 and - 0.77), meaning high correlation to cellular tissue region. For the mouse kidney, four features with high co-localization scores to extracellular space were selected: *m*/*z* 836.4425 with 0.79, *m*/*z* 1,105.5731 with 0.71, *m*/*z* 1,239.6418 with 0.71, and *m*/*z* 1,443.6943 with 0.75. Since the lowest co-localization score obtained for the mouse kidney was −0.55, no feature correlated with the cellular tissue region was selected in this case. Instead, feature *m*/*z* 1,198.7003 was selected due to its moderate correlation with the red subregion and its well documented identity in literature^8,36^. MS1 ion images of all selected precursors are shown in Figure 4B and an overview of the statistical analysis is given in Supplementary Table 4.

**Figure 4:**
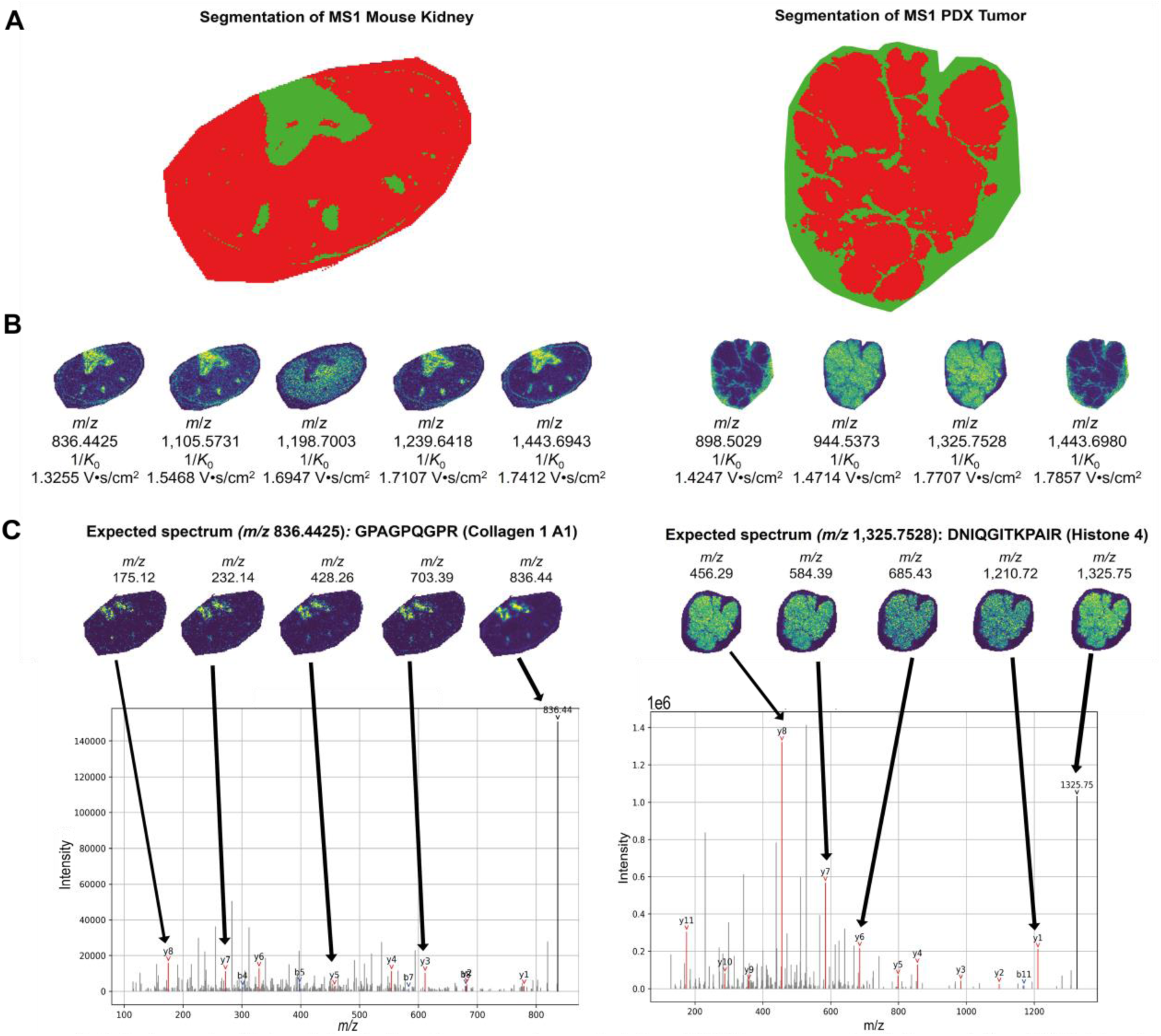
Selection and multiplexed identification of precursors in one single iprm-PASEF measurement **(A)** Segmentation of MSI datasets conducted in SCILS Lab are shown resulting in two subregions. Red subregion corresponds to the cellular space and the green subregion to the extracellular space. (B) MS1 ion images of precursors selected for iprm-PASEF. (C) MS2 spectra acquired in iprm-PASEF mode with a spatial resolution of 100 µm are shown for one selected precursor of each test tissue. B-ions are highlighted in red and y-ions in blue. Ion images of respective fragments are shown on top of each spectrum.

#### 3.5.3 iprm-PASEF

For iprm-PASEF of the mouse kidney, the set of five precursors chosen by exploratory statistical analysis was expanded with two randomly chosen, additional precursors due to 1/*K*_0_ window availability (Supplementary Table 5). The iprm-PASEF measurement of the PDX tumor employed a set of four predefined precursors (Supplementary Table 5).

The iprm-PASEF measurements generated 5,666 pixels in 47 minutes (mouse kidney) and 10,406 pixels in 81 minutes (PDX tumor) with a spatial resolution of 100×100 µm, resulting in a measurement speed of approximately 2 pixels per second which is less than 100 min per cm^2^ of tissue. Mean MS2 spectra for each precursor were separated by their 1/*K*_0_ traces and submitted to the search engine MASCOT, allowing for the assignment of peptide identities to the measured spectra. All precursors determined by MS1 statistical analysis were successfully identified by MASCOT (Table 2, Figure 4B). In contrast, the identity of the randomly selected precursors for the mouse kidney remained elusive. Mass accuracies were lower than 20 ppm for all precursors. The MASCOT score represents a statistical evaluation provided by Matrix Science; the higher the score, the more confidently a PSM can be considered^31^. The scores in this study ranged from 23.8 to 89.9 and were comparable with scores obtained in other published in situ MALDI MS/MS studies^37–40^. Presently, all acquired MS2 spectra per precursor over the entire tissue region are being averaged to yield a single MS2 spectrum, which we employed for the PSM step. We consider the possibility that quality filter steps prior to MS2 spectra averaging (e.g. signal-to-noise thresholds, subregion selection) may further improve final MS/MS data quality. However, this aspect was not further investigated in the present proof of concept study.

**Table 2:**
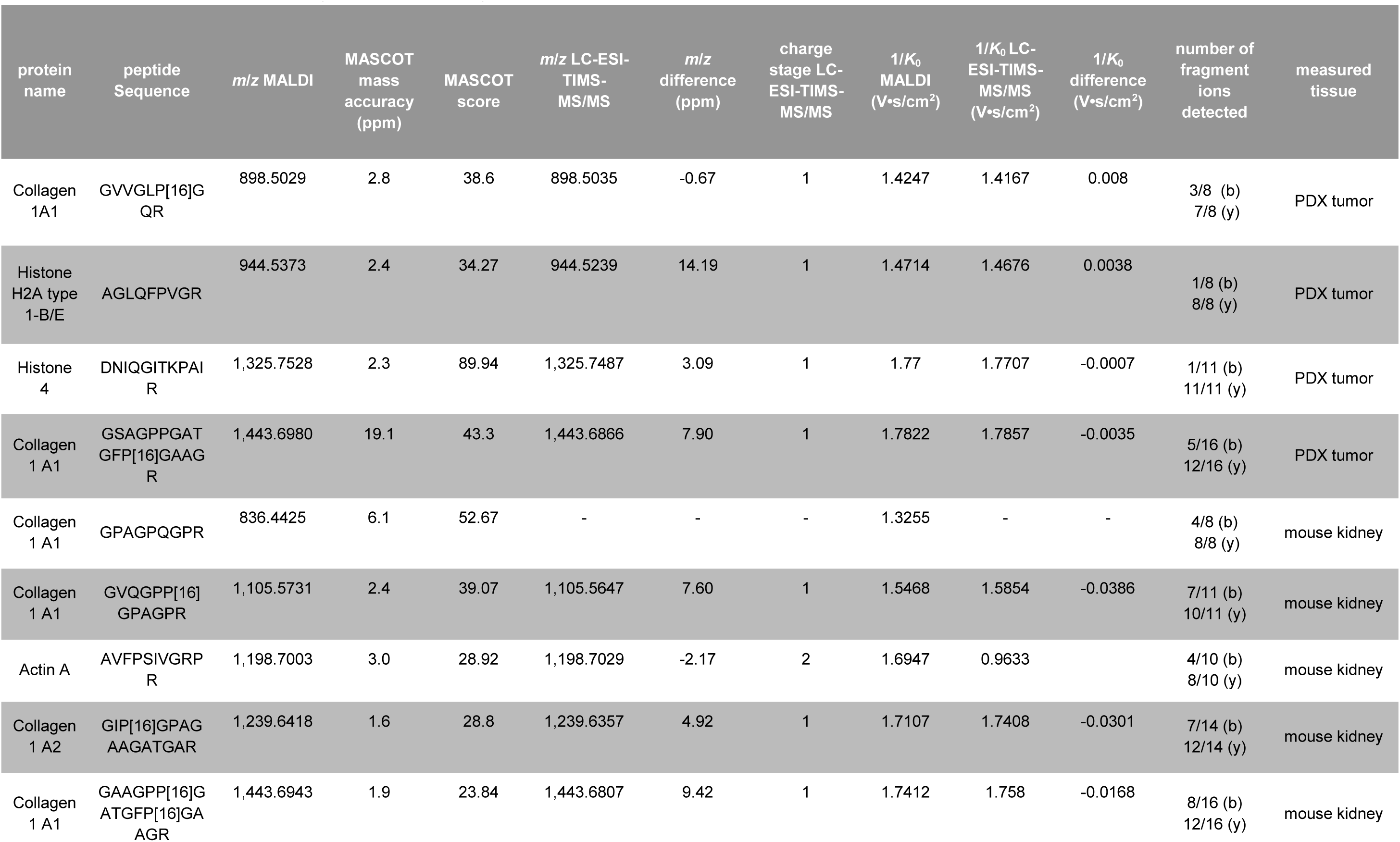
Overview of peptide identity corroborations by iprm-PASEF and LC-ESI-TIMS-MS/MS measurements of two test tissues.

#### 3.5.4 Corroboration Tools for Peptide Identification

To corroborate the peptide identifications obtained by MASCOT, we performed a number of additional analyses (Figure 1C): Since the online MASCOT tool only provides an identification score, the pyteomics package was employed to examine theoretical b- and y-ions of identified sequences as corroboration. All detected b- and y-ions and the mean MS2 spectra can be found in the supplementary information (Supplementary Figure 3, Supplementary Table 6) and in Table 2. For instance, all possible y-ions and 50 % of b-ions for peptide GPAGPQGPR (*m*/*z* 836.4425, mouse kidney) and all possible y-ions and 9.1 % of possible b-ions of the respective peptide DNIQGITKPAIR (*m*/*z* 1,325.7528, PDX tumor) were detected. Additionally, fragment co-localization was investigated in SCiLS Lab to include the acquired spatial distribution. The “Find Values Co-Localized to Feature” tool was used with the tissue region and the parent ion *m*/*z* as the feature to correlate to. For demonstration purposes, the co-localization analysis was only performed on one precursor for each tissue: *m*/*z* 836.4425 for the mouse kidney and *m*/*z* 1,325.7528 for the PDX tumor. Co-localization scores ranged from 0.50 to 0.90 (PDX tumor) and 0.40 to 0.56 (mouse kidney) (Supplementary Table 7). Peptides with a C-terminal arginine are usually well detected in MALDI-TOF devices, explaining the high amount of identified y-ions^41^. The MS2 spectra and ion images of the fragment signals, including the percentage of detected b- and y-ions, are visualized for the two exemplary precursors in Figure 4C. In addition, all precursor identifications are consistent with the expected histomorphological regions determined in the segmentation analysis, providing confidence at a biological level. For instance, *m*/*z* 1,325.7528, a peptide corresponding to the cellular protein histone 4, is found within in the red (cellular) subregion and *m*/*z* 836.4425 a peptide corresponding to the extracellular matrix protein collagen 1A1 is found within the green (extracellular) subregion.

To demonstrate further corroboration by LC-ESI-TIMS-MS/MS, we extracted the residual co-crystallized peptides after the MALDI imaging measurement and submitted them to LC-ESI-TIMS-MS/MS analysis^12^. By focusing on singly charged peptides, we detected seven out of nine precursors in 1+ charge, allowing for the comparison of 1/*K*_0_ in addition to the *m*/*z* value. One precursor (*m*/*z* 1,198.7003) was only detected on 2+ charge and one precursor at *m*/*z* 836.4425 failed to be detected in LC-ESI-TIMS-MS/MS. For the mouse kidney, *m*/*z* variance was lower than 10 ppm and 1/*K*_0_ variance lower than 0.04 V•s/cm^2^. For the PDX tumor, the variance for *m*/*z* was lower than 15 ppm and for 1/*K*_0_ lower than 0.01 V•s/cm^2^ (Table 2).

Using these two test FFPE tissues, we demonstrated that iprm-PASEF enables multiplexed in situ MS/MS, making it a valuable tool in the field of spatial proteomics.

## 4 Discussion

In this study, we present the first performance of iprm-PASEF on tryptic peptides as an initial step towards improving MALDI-based spatial proteomics. This method enables us to tackle the issue of efficient spatial peptide identification as a key challenge in MALDI imaging of peptides. We demonstrate the ability to identify peptidic precursor ions on two different FFPE tissue samples in a spatially resolved manner using iprm-PASEF. Peptide identification was corroborated using the MASCOT PSM search of MS2 spectra and additional LC-ESI-TIMS-MS/MS measurements. For this proof of concept, we targeted frequently identified and highly abundant features that were only multiplexed up to five precursors. The ability to identify low abundant features remains to be investigated, as does the possible number of precursors identified per run. Nevertheless, identifying high abundant features remains relevant as they may also play crucial roles in biological contexts, such as collagens in the tumor microenvironment and cancer pathogenesis^42–47^. In addition, the identification of frequently reported features remains desirable in each individual experimental sample, as MS1 matching can yield multiple putative peptide identifications and vary between tissue specimens.

Notably, the presented technique is highly adaptable and can be customized to suit specific experimental needs. Several potential improvements could be tackled: First, MS1 imaging and iprm-PASEF could be conducted on the same slide with a laser offset, reducing sample usage and simplifying co-registration. Using a laser offset could also allow for multiple iprm-PASEF runs with different precursor lists on the same slide, thereby increasing the number of targeted precursors. Second, the iprm-PASEF data analysis pipeline presented in this study relies on four different software programs (DataAnalysis, SCiLS Lab, MASCOT, Python/pyteomics), requiring multiple import and export steps. Manual adjustments were necessary due to e.g. limited performance of the SCiLS Lab feature detection tool for ion mobility data. This highlights potential for further optimization and automation of the data analysis workflow.

As a targeted method, iprm-PASEF is restricted to a defined number of precursors and does not represent an exploratory approach. This targeted nature implements the precursor selection as a highly adjustable step in the presented pipeline that is dependent on prior information. However, it can be individually tailored to a scientific goal. Given that this study is a technical proof of concept, we focused on exploratory statistical analyses of the MALDI MS1 imaging data including segmentation and co-localization for the precursor selection. Other approaches for manual precursor selection may include filtering the MS1 data for the most intense signals, literature-based searches for specific peptides/proteins of interest, in silico digestion of specific proteins of interest, and generating LC-ESI-TIMS-MS/MS-based libraries from measurements of the same or comparable tissue sample. In contrast, Spatial Ion Mobility Scheduled Exhaustive Fragmentation (SIMSEF), a tool based on optimized and automated scheduling of MS/MS events, offers potential for non-targeted MS2 bioimaging^48^. However, its applicability is currently limited to metabolites and would require major technical adjustments to make it suitable for tryptic peptides. In addition, the SIMSEF tool would not provide spatial fragment ion information to confirm correct peptide identification at each pixel and would require major advances in data analysis.

Compared to other spatial proteomics methods, the presented workflow is not dependent on antibody availability, specificity or individual sample preparation establishment. This is the case in other multiplexed antibody-based methodologies such as CODEX (Akoya), Imaging Mass Cytometry, multiplex immunofluorescence/immunohistochemistry (IHC) or MALDI-IHC^49–52^. Another powerful and upcoming method is MS-based deep visual proteomics that exploits laser microdissection and provides a high spatial proteome coverage. However, compared to MALDI imaging it is extremely laborious and requires a high amount of measurement and sample preparation time, making routine usage challenging^53,54^.

Overall, by integrating TIMS and PASEF into MALDI imaging, we established a spatial proteomics workflow that enables multiplexed peptide identification, is low-cost, fast, independent of antibodies and easy to combine with other analysis modalities. Altogether, this pipeline includes several tools for corroboration to increase confidence in peptide identification in MALDI imaging, making it highly valuable as a first MALDI-based and straight forward approach in the field of spatial proteomics.

## 5 Author Contributions

Mujia Jenny Li: conceptualization, data curation, formal analysis, methodology, writing – original draft. Larissa Chiara Meyer: conceptualization, data curation, formal analysis, methodology, writing – original draft. Nadine Meier: data curation. Jannik Witte: conceptualization, software. Maximilian Maldacker: methodology. Adrianna Seredynska: methodology. Julia Schueler: resources. Oliver Schilling: conceptualization, project administration, funding acquisition, supervision, writing – review and editing. Melanie Christine Föll: conceptualization, project administration, funding acquisition, supervision, writing – review and editing.

## Acknowledgements

The authors thank Prof. Dr. Thomas Reinheckel and Dr. Martina Tholen including their laboratory for providing tissue samples. Further, we want to thank Dr. Andras Kiss and Dr. Arne Behrens from Bruker Daltonics for their technical support.

## 6 Funding Statement

Prof. Dr. Oliver Schilling acknowledges funding by the Deutsche Forschungsgemeinschaft (DFG, projects 446058856, 466359513, 444936968,

405351425, 431336276, 43198400 (SFB 1453 “NephGen”), 441891347 (SFB 1479 “OncoEscape”), 423813989 (GRK 2606 “ProtPath”), 322977937 (GRK 2344 “MeInBio”), the ERA PerMed program (BMBF, 01KU1916, 01KU1915A), the ERA TransCan program (BMBF 01KT2201,“PREDICO”; 01KT2333, “ICC-STRAT”), the German Consortium for Translational Cancer Research (project Impro-Rec), the investBW program (project BW1_1198/03 “KASPAR”), the BMBF KMUi program (project 13GW0603E, project ESTHER), BMBF FKZ 03ZU1208AA (nanodiag BW).

## 7 Data Availability

The raw data is publicly available on MassIVE (Mass Spectrometry Interactive Virtual Environment) under the following links:

- Kidney MS1 data: doi:10.25345/C54Q7R252 (MSV000096373)
- Kidney MS/MS data: doi:10.25345/C5W669M54 (MSV000096375)
- PDX MS1 data: doi:10.25345/C50Z71819 (MSV000096374)
- PDX MS/MS data: doi:10.25345/C5RF5KT0H (MSV000096376)

## Abbreviations

ACN: Acetonitrile
AmBC: Ammonium bicarbonate
CHCA: Alpha-cyano-4-hydroxycinnamic acid matrix
DDA: Data-dependent acquisition
ESI: Electrospray ionization
FFPE: Formalin-fixed paraffin-embedded
IHC: Immunohistochemistry
iprm-PASEF: Imaging Parallel Reaction Monitoring, Parallel Accumulation SErial Fragmentation
ITO: Indium tin oxide
LC-MS/MS: Liquid-chromatography tandem mass spectrometry
m/z: Mass over charge
MALDI: Matrix-assisted laser desorption/ionization
MS: Mass spectrometry
prm: Parallel Reaction Monitoring
PASEF: Parallel Accumulation SErial Fragmentation
PDX: Patient-derived xenografts
PSM: Peptide to spectrum matching
Q-TOF: Quadrupole time-of-flight
RMS: Root mean square
TFA: Trifluoroacetic acid
TIMS: Trapped Ion Mobility Spectrometry
1/*K*_0_: Inverse of reduced ion mobility

