## supplementary tables for "Spatial Proteomics By Parallel Accumulation-Serial Fragmentation Supported MALDI MS/MS Imaging: A First Glance Into Multiplexed and Spatial Peptide Identification"

**Supplementary Table 1**: Precursors selected for testing effects of including TIMS dimension on MS/MS level.

| ***m*/*z*** | **1/*K*_0_ (V•s/cm^2^)** | **Peptide_genename** |
| --- | --- | --- |
| 944.53 | 1.472 | AGLQFPVGR_H2A |
| 1,198.70 | 1.675 | AVFPSIVGRPR_ACTA |
| 1,325.75 | 1.750 | DNIQGITKPAIR_H4 |

**Supplementary Table 2: Number of b- and y-ions detected in iprm-PASEF measurements of 5 synthetic peptides.** A mastermix of 5 synthetic peptides was spotted onto an ITO slide. Precursors were either targeted one after another in an iprm-PASEF 1-plex setup or all together in an iprm-PASEF 5-plex setup. The fragmentation pattern including series of detected b- and y-ions for each of the targeted peptides is shown in this table.

| **Peptide sequence** | ***m/z* MALDI** | **1/*K*_0_ start** | **1/*K*_0_ end** | **Number of fragment ions detected** | **Detected b-ions** | **Detected y-ions** | **Experiment** |
| --- | --- | --- | --- | --- | --- | --- | --- |
| LGGNEQVTR | 973.5061 | 1.41 | 1.47 | 1/8 (b) 8/8 (y) | *m/z* (b₃)  246.18 | *m/z* (y_1_) 860.42  *m/z* (y₂) 803.40  *m/z* (y₃) 746.38  *m/z* (y₄) 632.33  *m/z* (y₅) 503.29  *m/z* (y₆) 375.23  *m/z* (y₇) 276.17  *m/z* (y₈) 175.12 | 1-plex iprm-PASEF |
| RPKPQQFFGLM | 1347.73 | 1.75 | 1.79 | 4/10 (b) 3/10 (y) | *m/z* (b_1_)  175.12  *m/z* (b_3_)  400.27  *m/z* (b_4_)  497.32  *m/z* (b₆)  753.43 | *m/z* (y₆) 614.26  *m/z* (y₇) 467.19  *m/z* (y_11_) 320.21 | 1-plex iprm-PASEF |
| QRPRLSHKGPMPF | 1550.84 | 1.797 | 1.83 | 4/12 (b) 6/12 (y) | *m/z* (b_4_)  556.33  *m/z* (b_5_)  669.42  *m/z* (b_6_)  756.45  *m/z* (b_12_)  1403.76 | *m/z* (y₂) 1266.67  *m/z* (y₅) 900.43  *m/z* (y₆) 813.40  *m/z* (y₇) 676.35  *m/z* (y₈) 548.25  *m/z* (y_11_) 263.14 | 1-plex iprm-PASEF |
| KLKESYCQRQGVPMN | 1780.8835 | 1.87 | 1.93 | 7/14 (b) 10/14 (y) | *m/z* (b_3_)  388.23  *m/z* (b_4_)  517.33  *m/z* (b_5_)  604.30  *m/z* (b_6_)  767.43  *m/z* (b_8_)  998.42  *m/z* (b_10_)  1282.56  *m/z* (b_14_)  1666.84 | *m/z* (y_4_) 1282.56  *m/z* (y_5_) 1195.53  *m/z* (y_6_) 1032.47  *m/z* (y_7_) 929.46  *m/z* (y_9_) 801.40  *m/z* (y_9_) 645.30  *m/z* (y_10_) 517.19  *m/z* (y_11_) 460.22  *m/z* (y_12_) 361.15  *m/z* (y_13_) 264.10 | 1-plex iprm-PASEF |
| KLKVIGQDSSEIHFKV | 1828.0327 | 1.97 | 2.1 | 6/15 (b) 10/15 (y) | *m/z* (b_3_)  388.23  *m/z* (b_5_)  600.30  *m/z* (b_6_)  657.44  *m/z* (b7)  785.52  *m/z* (b_8_)  900.55  *m/z* (b_9_)  987.58 | *m/z* (y_4_) 1359.68  *m/z* (y_5_) 1246.60  *m/z* (y_7_) 1061.52  *m/z* (y_8_) 946.50  *m/z* (y_9_) 859.46  *m/z* (y_10_) 772.43  *m/z* (y_11_) 643.39  *m/z* (y_12_) 530.31  *m/z* (y_13_) 393.25  *m/z* (y_14_) 246.18 | 1-plex iprm-PASEF |
| LGGNEQVTR | 973.5061 | 1.41 | 1.47 | 1/8 (b) 8/8 (y) | *m/z* (b₃)  246.18 | *m/z* (y_1_) 860.42  *m/z* (y₂) 803.40  *m/z* (y₃) 746.38  *m/z* (y₄) 632.33  *m/z* (y₅) 503.29  *m/z* (y₆) 375.23  *m/z* (y₇) 276.17  *m/z* (y₈) 175.12 | 5-plex iprm-PASEF |
| RPKPQQFFGLM | 1347.73 | 1.75 | 1.79 | 5/10 (b) 3/10 (y) | *m/z* (b_1_)  175.12  *m/z* (b_3_)  400.27  *m/z* (b_4_)  497.32  *m/z* (b₆)  753.43  *m/z* (b_7_)  900.49 | *m/z* (y₆) 614.26  *m/z* (y₇) 467.19  *m/z* (y₈) 320.21 | 5-plex iprm-PASEF |
| QRPRLSHKGPMPF | 1550.84 | 1.797 | 1.83 | 5/12 (b) 8/12 (y) | *m/z* (b_3_)  400.23  *m/z* (b_4_)  556.33  *m/z* (b_5_)  669.42  *m/z* (b_6_)  756.45  *m/z* (b_12_)  1403.76 | *m/z* (y₂) 1266.67  *m/z* (y₄) 1013.52  *m/z* (y₅) 900.43  *m/z* (y₆) 813.40  *m/z* (y₇) 676.35  *m/z* (y₈) 548.25  *m/z* (y_9_) 491.23  *m/z* (y_11_) 263.14 | 5-plex iprm-PASEF |
| KLKESYCQRQGVPMN | 1780.8835 | 1.87 | 1.93 | 7/14 (b) 10/14 (y) | *m/z* (b_3_)  388.23  *m/z* (b_4_)  517.33  *m/z* (b_5_)  604.30  *m/z* (b_6_)  767.43  *m/z* (b_8_)  998.42  *m/z* (b_10_)  1282.56  *m/z* (b_14_)  1666.84 | *m/z* (y_4_) 1282.56  *m/z* (y_5_) 1195.53  *m/z* (y_6_) 1032.47  *m/z* (y_7_) 929.46  *m/z* (y_8_) 801.40  *m/z* (y_9_) 645.30  *m/z* (y_10_) 517.19  *m/z* (y_11_) 460.22  *m/z* (y_12_) 361.15  *m/z* (y_13_) 264.10 | 5-plex iprm-PASEF |
| KLKVIGQDSSEIHFKV | 1828.0327 | 1.97 | 2.1 | 7/15 (b) 11/15 (y) | *m/z* (b_3_)  388.23  *m/z* (b_4_)  487.21  *m/z* (b_5_)  600.30  *m/z* (b_6_)  657.44  *m/z* (b_7_)  785.52  *m/z* (b_8_)  900.55  *m/z* (b_9_)  987.58 | *m/z* (y_4_) 1359.68  *m/z* (y_5_) 1246.60  *m/z* (y_6_) 1189.58  *m/z* (y_7_) 1061.52  *m/z* (y_8_) 946.50  *m/z* (y_9_) 859.46  *m/z* (y_10_) 772.43  *m/z* (y_11_) 643.39  *m/z* (y_12_) 530.31  *m/z* (y_13_) 393.25  *m/z* (y_14_) 246.18 | 5-plex iprm-PASEF |

**Supplementary Table 3**: Optimization of the LC-ESI-TIMS-MS/MS measurement focused on singly charged peptides. 100 ng HeLa digests were measured in three different measurement settings for optimization of 1+ peptide detection. Number of +1 peptide identifications with “Match-type”= MS/MS.

|  | **Prepulse storage** | **Transfer time** | **Def Delta 1** | **Funnel 1 RF** | **Funnel 2 RF** | **Collision energy** | **Collision RF** | **Ion Energy** | **1/*K*_0_ range** | **TIMS in pressure** | **+1 Peptide ID** |
| --- | --- | --- | --- | --- | --- | --- | --- | --- | --- | --- | --- |
| DDAPasef Proteomics | 12 µs | 60 µs | 70 V | 450 Vpp | 200 Vpp | 20-59 eV | 1,500 Vpp | 5eV | 0.6-1.6 V*s/cm^2^ | 2.4 mbar | 0 |
| DDAPasef 1+ | 15 µs | 120 µs | 70 V | 450 Vpp | 200 Vpp | 30-74 eV | 2,000 Vpp | 5eV | 0.7-1.75 V*s/cm^2^ | 2.4 mbar | 2,359 |
| DDAPasef 1+ | 15 µs | 120 µs | 70 V | 450 Vpp | 200 Vpp | 30-74 eV | 2,000 Vpp | 5eV | 0.7-2.1 V*s/cm^2^ | 2 mbar | 3,211 |
| DDAPasef 1+ opt | 15 µs | 70 µs | 80 V | 250 Vpp | 500 Vpp | 20-75 eV | 2,500 Vpp | 5eV | 0.7-2.1 V*s/cm^2^ | 2 mbar | 3,646 |

**Supplementary Table 4: Co-localization scores for statistical analysis.** Overview of features chosen for the final precursor list with a significant co-localization scores (CS) determined by the “Find Values Co-Localized to Region” tool in SCiLS Lab. For *m*/*z* 944.5373, 1,325.7528, 1,443.698, 836.4425, 1,198.7003, 1,105.5731 and 1,443.6943, literature was found providing putative peptide identifications and citations are included in the table.

| ***m*/*z*** | **1/*K*_0_ (V•s/cm^2^)** | **CS** | **Tissue** | **Literature** |
| --- | --- | --- | --- | --- |
| 898.5029 | 1.4247 | 0.74 | PDX tumor | - |
| 944.5373 | 1.4714 | -0.71 | PDX tumor | PMID: 27061135 |
| 1,325.7528 | 1.77 | -0.77 | PDX tumor | PMID: 34206844, 31664609,34572274,38492056,18712763,38928454 |
| 1,443.698 | 1.7822 | 0.76 | PDX tumor | PMID: 26505774 |
| 836.4425 | 1.32 | 0.79 | mouse kidney | PMID: 27696080, 30548962, 36768889, 34572274, 31664609, 27939604 |
| 1,198.7003 | 1.69 | -0.15 | mouse kidney | PMID: 35684402, 27939604 |
| 1,105.5731 | 1.55 | 0.74 | mouse kidney | PMID: 27061135 |
| 1,239.6418 | 1.71 | 0.71 | mouse kidney | - |
| 1,443.6943 | 1.74 | 0.75 | mouse kidney | PMID: 26505774 |

**Supplementary Table 5:** All precursor lists that were submitted to iprm-PASEF analysis.

**A:** Mouse kidney precursor list for iprm-PASEF. Precursors 1260.61 *m*/*z* and 852.43 *m*/*z* were included because of ion mobility window availability but could not be identified by MASCOT.

| ***m*/*z*** | **1/*K*_0_ start** | **1/*K*_0_ end** |
| --- | --- | --- |
| 836.43 | 1.31 | 1.35 |
| 852.43 | 1.27 | 1.3 |
| 1,106.58 | 1.44 | 1.48 |
| 1,199.7 | 1.68 | 1.7 |
| 1,239.64 | 1.701 | 1.73 |
| 1,260.61 | 1.64 | 1.67 |
| 1,443.6819 | 1.7301 | 1.78 |

**B:** PDX tumor precursor list for iprm-PASEF

| ***m*/*z*** | **1/*K*_0_ start** | **1/*K*_0_ end** |
| --- | --- | --- |
| 898.5 | 1.4 | 1.43 |
| 944.5 | 1.441 | 1.49 |
| 1,325.72 | 1.75 | 1.775 |
| 1,443.67 | 1.781 | 1.81 |

**Supplementary Table 6:** All precursors targeted with iprm-PASEF including the detected b- and y-ions calculated using the pyteomics package in python.

| **Peptide sequence** | ***m/z* MALDI** | **Detected b-ions** | **Detected y-ions** | **Measured tissue** |
| --- | --- | --- | --- | --- |
| GVVGLP[16]GQR | 898.5029 | *m/z* (b₂) 175.12  *m/z* (b₃) 274.19  *m/z* (b₈) 742.42 | *m/z* (y₂) 742.42  *m/z* (y₃) 643.35  *m/z* (y₄) 586.33  *m/z* (y₅) 473.25  *m/z* (y₆) 360.20  *m/z* (y₇) 303.18  *m/z* (y₈) 175.12 | PDX tumor |
| AGLQFPVGR | 944.5373 | *m/z* (b₈) 788.42 | *m/z* (y₁) 873.49  *m/z* (y₂) 816.47  *m/z* (y₃) 703.39  *m/z* (y₄) 575.33  *m/z* (y₅) 428.26  *m/z* (y₆) 331.21  *m/z* (y₇) 232.14  *m/z* (y₈) 175.12 | PDX tumor |
| DNIQGITKPAIR | 1325.7528 | *m/z* (b₁₁) 1169.65 | *m/z* (y₁) 1210.72  *m/z* (y₂) 1096.68  *m/z* (y₃) 983.60  *m/z* (y₄) 855.54  *m/z* (y₅) 798.52  *m/z* (y₆) 685.43  *m/z* (y₇) 584.39  *m/z* (y₈) 456.29  *m/z* (y₉) 359.24  *m/z* (y₁₀) 288.20  *m/z* (y₁₁) 175.12 | PDX tumor |
| GSAGPPGATGFP[16]GAAGR | 1443.698 | *m/z* (b₅) 388.18  *m/z* (b₆) 485.22  *m/z* (b₇) 542.26  *m/z* (b₈)613.32  *m/z* (b₁₂) 1031.47 | *m/z* (y₂) 1299.61  *m/z* (y₆) 977.48  *m/z* (y₇) 920.46  *m/z* (y₈) 849.42  *m/z* (y₉) 748.37  *m/z* (y₁₀) 691.35  *m/z* (y₁₁) 544.28  *m/z* (y₁₂) 431.23  *m/z* (y₁₃) 374.21  *m/z* (y₁₄) 303.18  *m/z* (y₁₅) 232.14  *m/z* (y₁₆) 175.12 | PDX tumor |
| GPAGPQGPR | 836.4425 | *m/z* (b₄) 301.15  *m/z* (b₅) 398.20  *m/z* (b₇) 583.28  *m/z* (b₈)680.34 | *m/z* (y₁) 779.41  *m/z* (y₂) 682.36  *m/z* (y₃) 611.33  *m/z* (y₄) 554.30  *m/z* (y₅) 457.25  *m/z* (y₆) 329.19  *m/z* (y₇) 272.17  *m/z* (y₈) 175.12 | mouse kidney |
| GVQGPP[16]GPAGPR | 1105.5731 | *m/z* (b₂) 175.12  *m/z* (b₃) 303.03  *m/z* (b₇) 627.21  *m/z* (b₈) 724.33  *m/z* (b₉) 795.30  *m/z* (b₁₀) 852.39  *m/z* (b₁₁) 949.48 | *m/z* (y₂) 949.48  *m/z* (y₃) 821.42  *m/z* (y₄) 764.40  *m/z* (y₅) 667.35  *m/z* (y₆) 554.30  *m/z* (y₇) 497.28  *m/z* (y₈) 400.23  *m/z* (y₉) 329.19  *m/z* (y₁₀) 272.17  *m/z* (y₁₁) 175.12 | mouse kidney |
| AVFPSIVGRPR | 1198.7003 | *m/z* (b₆) 633.32  *m/z* (b₇) 732.41  *m/z* (b₉) 945.55  *m/z* (b₁₀) 1042.60 | *m/z* (y₃) 881.53  *m/z* (y₄) 784.48  *m/z* (y₅) 697.45  *m/z* (y₆) 584.36  *m/z* (y₇) 485.29  *m/z* (y₈) 428.27  *m/z* (y₉) 272.17  *m/z* (y₁₀) 175.12 | mouse kidney |
| GIP[16]GPAGAAGATGAR | 1239.6418 | *m/z* (b₄) 359.13  *m/z* (b₅) 456.22  *m/z* (b₆) 527.25  *m/z* (b₇) 584.31  *m/z* (b₈) 655.38  *m/z* (b₉) 726.32  *m/z* (b₁₀) 783.36 | *m/z* (y₂) 1069.54  *m/z* (y₄) 899.47  *m/z* (y₅) 802.41  *m/z* (y₆) 731.38  *m/z* (y₇) 674.36  *m/z* (y₈) 603.32  *m/z* (y₉) 532.28  *m/z* (y₁₀) 475.26  *m/z* (y₁₁) 404.22  *m/z* (y₁₂) 303.18  *m/z* (y₁₃) 246.16  *m/z* (y₁₄) 175.12 | mouse kidney |
| GAAGPP[16]GATGFP[16]GAAGR | 1443.6943 | *m/z* (b₅) 372.07  *m/z* (b₆) 485.15  *m/z* (b₇) 542.21  *m/z* (b₈) 613.25  *m/z* (b₉) 714.31  *m/z* (b₁₀) 771.28  *m/z* (b₁₁) 918.39  *m/z* (b₁₂) 1031.36 | *m/z* (y₄) 1187.58  *m/z* (y₆) 977.48  *m/z* (y₇) 920.46  *m/z* (y₈) 849.42  *m/z* (y₉) 748.37  *m/z* (y₁₀) 691.35  *m/z* (y₁₁) 544.28  *m/z* (y₁₂) 431.23  *m/z* (y₁₃) 374.21  *m/z* (y₁₄) 303.18  *m/z* (y₁₅) 232.14  *m/z* (y₁₆) 175.12 | mouse kidney |

**Supplementary Table 7:** Co-localization scores from fragment ions for corroboration of iprm-PASEF peptide identification. Generated using “Find Values Co-Localized to Feature” tool in SCiLS Lab.

| ***m*/*z* precursor** | **Fragment ion** | **Co-localization score** | **Tissue** |
| --- | --- | --- | --- |
| 836.4425 | 611.328 (y_3_) | 0.4599 | mouse kidney |
| 836.4425 | 554.3025 (y_4_) | 0.4949 | mouse kidney |
| 836.4425 | 397.2158 (b_5_) | 0.5625 | mouse kidney |
| 836.4425 | 329.1929 (y_6_) | 0.4699 | mouse kidney |
| 836.4425 | 272.1689 (y_7_) | 0.4035 | mouse kidney |
| 1,325.7528 | 1,210.7247 (y_1_) | 0.9683 | PDX tumor |
| 1,325.7528 | 855.5296 (y_4_) | 0.7638 | PDX tumor |
| 1,325.7528 | 798.5168 (y_5_) | 0.6254 | PDX tumor |
| 1,325.7528 | 685.4353 (y_6_) | 0.5655 | PDX tumor |
| 1,325.7528 | 584.3861 (y_7_) | 0.7317 | PDX tumor |
