## Supplementary figures and images for "Spatial Proteomics By Parallel Accumulation-Serial Fragmentation Supported MALDI MS/MS Imaging: A First Glance Into Multiplexed and Spatial Peptide Identification"

### supplementary figures

#### Slide 1
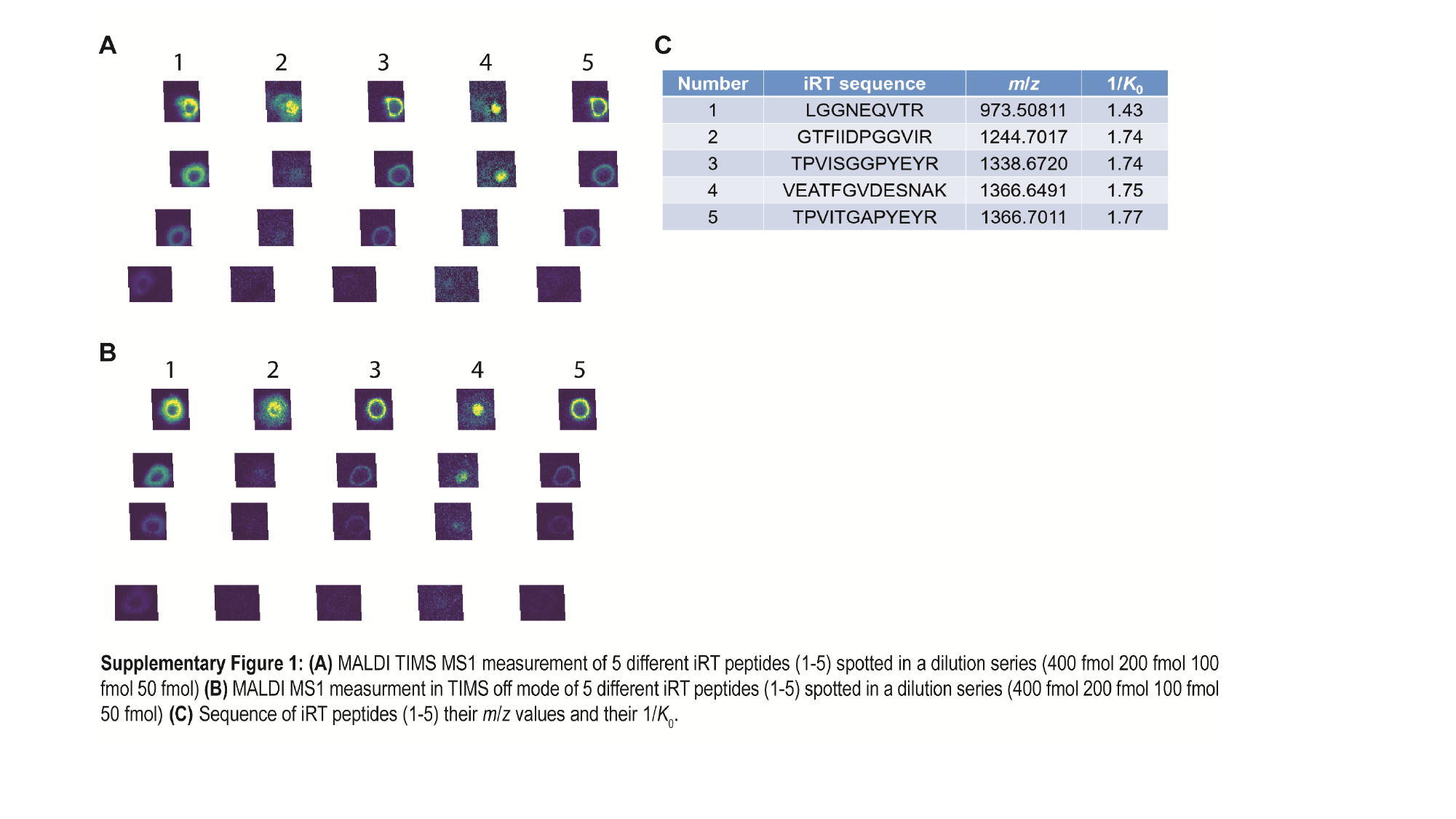

#### Slide 2
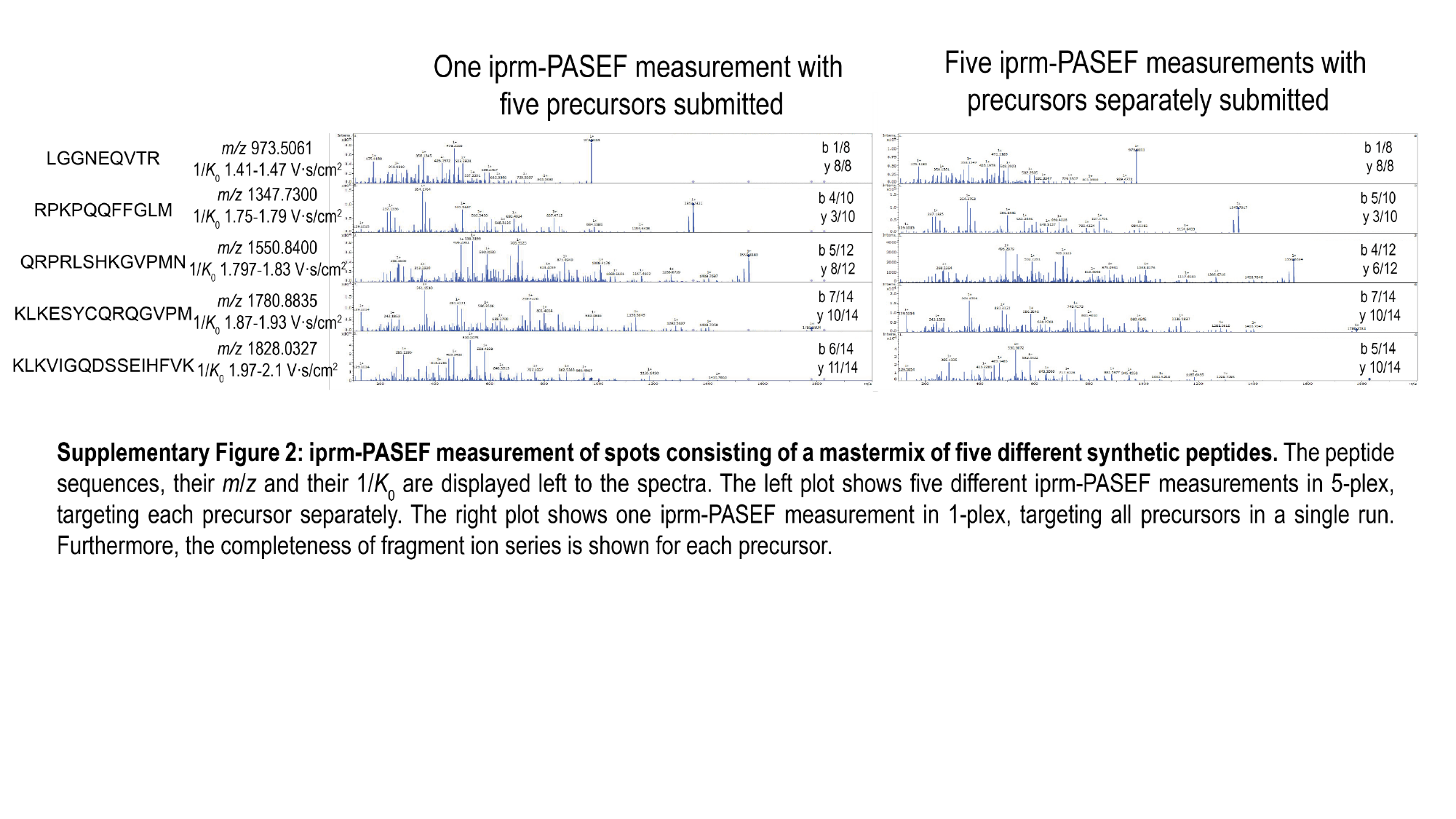

#### Slide 3
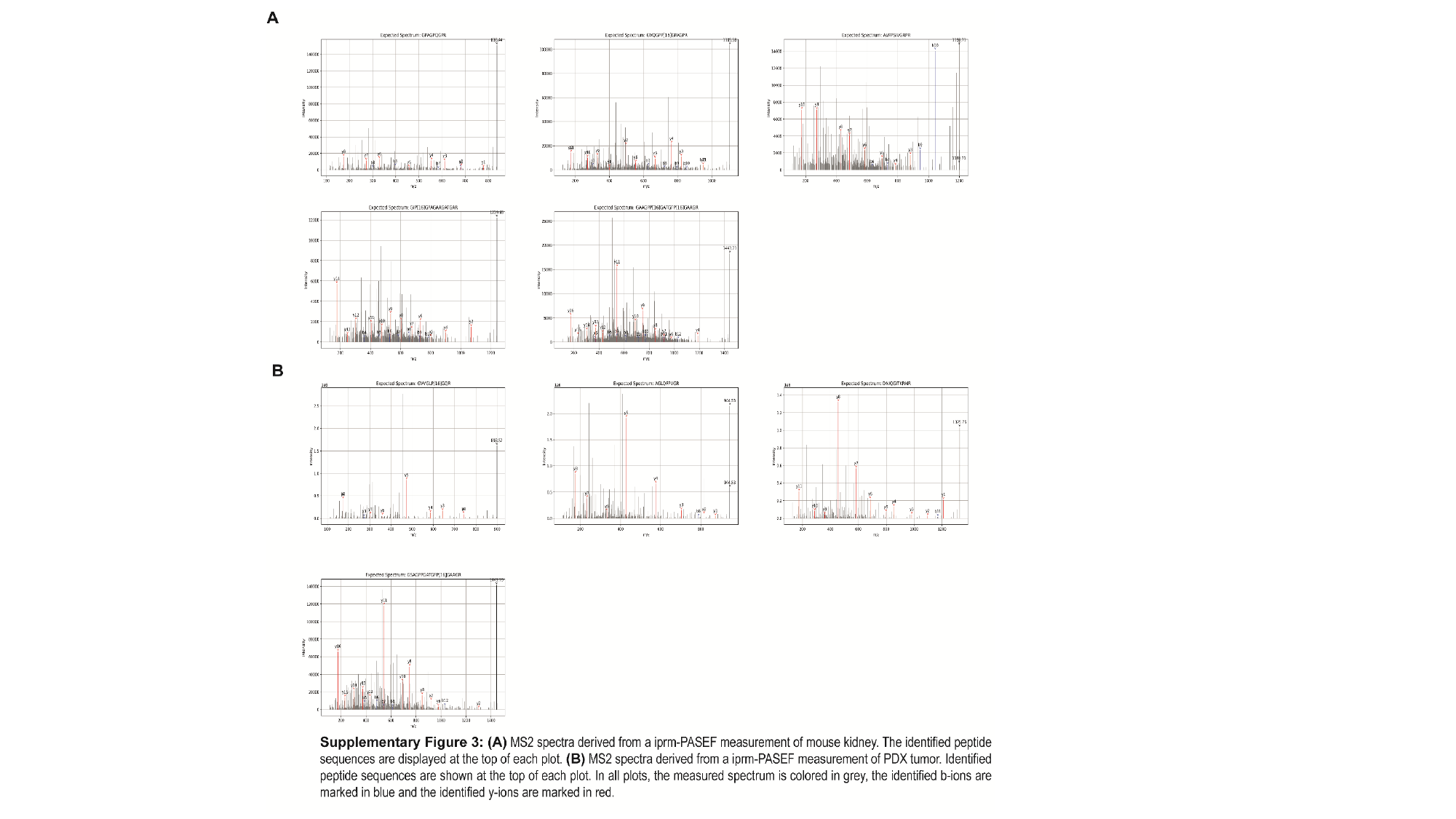
